# An iterative design approach to development of an ex-vivo normothermic multivisceral perfusion platform

**DOI:** 10.1101/2024.10.04.616696

**Authors:** L. Leonie van Leeuwen, Matthew L. Holzner, Ceilidh McKenney, Rachel Todd, Jamie K. Frost, S. Gudibendi, Leona Kim-Schluger, Thomas Schiano, Sander Florman, M. Zeeshan Akhtar

## Abstract

Challenges in normothermic machine perfusion (NMP) remain, particularly concerning the duration for which individual organs can be safely preserved. We hypothesize that optimal preservation can be achieved by perfusing organs together in a multivisceral block. Therefore, our aim was to establish a platform for *ex vivo* multivisceral organ perfusion.

Multivisceral grafts containing the liver, kidneys, pancreas, spleen and intestine were obtained from Yorkshire pigs. Three generation (gen) setups were tested during the iterative design process, and minor changes were made throughout. Gen1 (n=4) used a custom-designed single perfusion circuit. Gen2 (n=3) employed a dual perfusion circuit. Gen3 (n=4) featured a single perfusion circuit with an optimized basin and reservoir. Grafts underwent NMP using an autologous blood-based perfusate, while hemostatic parameters and function were assessed.

With each iteration, aortic flow improved, resistance decreased, urine output increased, oxygen consumption rose, perfusate lactate levels dropped, and pH stability improved. Cellular injury trended lower in Gen3. Histological evaluation demonstrated minimal differences in Gen2 and 3.

We demonstrate the feasibility of abdominal multivisceral NMP for up to 8 hours. Adequate arterial flow, stable perfusate pH, and high oxygen consumption in setup 3 indicate organ viability. Multivisceral perfusion may serve as a platform for long-term NMP.

## Introduction

The introduction of normothermic machine perfusion (NMP) has been a significant advancement in transplantation, affecting nearly all organs including heart, lung, liver, and kidney transplantation. This technology has rapidly accelerated the field, safely expanding the ability for *ex vivo* organ preservation, and allowing for the assessment, optimization, and repair of organs prior to transplantation.^1–6^ Used in combination with other preservation technologies, such as normothermic regional perfusion and hypothermic machine perfusion (HMP), NMP is expanding the boundaries of what are considered suitable organs for transplant.

Despite the advancements in machine preservation, several limitations persist, particularly concerning the duration for which individual organs can be safely preserved. While ongoing efforts aim to extend these boundaries, achieving near physiological conditions remains essential for advancing safe *ex vivo* storage. Isolated experimental liver perfusion beyond 24-36 hours often necessitates the use of dialysate filters, or entire dialysis circuits, to remove waste products not cleared by the liver itself to sustain prolonged perfusion.^7^

Preserving certain isolated organ systems, such as the kidney, pancreas, and small bowel, under normothermic conditions, even for short periods, continues to be challenging. In the case of kidneys, the rationale for choosing normothermic over hypothermic perfusion is still unclear, and the organ’s response to metabolic demands during isolated normothermic perfusion differs significantly from both hypothermic and *in situ* conditions as well as per device and protocol used.^8,9^ The pancreas, being a particularly susceptible low flow organ, is prone to substantial edema and injury during normothermic perfusion, making it difficult to maintain without damage.^10^ Similarly, normothermic perfusion of the bowel is challenging, with only a few groups successfully achieving it up to six hours.^11,12^ As a result, static cold storage or HMP under low flow states remains a simpler and more reliable option for some organ systems and under certain circumstances. However, normothermic environments offer the most suitable conditions for detailed biological assessments and the implementation of therapies, including cellular and regenerative treatments.

We hypothesize that maintaining organs *en bloc* may overcome the limitations of isolated organ perfusion by better preserving physiological *in situ* conditions and allow for viable long-term NMP. In this approach, multiple organs are recovered *en bloc*, connected within a single circuit, and maintained together throughout perfusion. The interactions and interplay between organ systems are likely critical for enabling long-term storage *ex vivo*. Achieving this goal could not only facilitate the selective and sequential harvest of organs for transplantation but also provide a valuable model for drug discovery and testing. Additionally, this approach could offer significant benefits to recipients of multi-visceral transplants, who are among the most challenging patients to manage in the field of transplantation.

Research on multivisceral perfusion is still limited.^13–16^ Therefore, the goal in this proof-of-concept study was to develop a multi-visceral perfusion platform using a rapid-prototyping and iterative design approach. The aim of this paper was to characterize perfusion dynamics, physiological parameters, and basic molecular and histological data from this iterative design process. Initial success was deemed to be the provision of steady state perfusion for at least eight hours.

## Materials and methods

### Rationale for Iterative Design Process

An iterative design approach was adopted for developing the multi-visceral perfusion platform, allowing continuous refinement through cycles of design, testing, and review. Significant modifications were made with each cycle, enabling early identification of issues and reducing the number of animals used. This method allowed for targeted improvements based on real-time experimental findings, enhancing the platform’s functionality and reliability.

### Generation 1 Perfusion Set-Up (gen 1)

The first-generation platform featured a custom-built open perfusion circuit designed to maintain continuous flow and oxygenation to the organs. The circuit consisted of a centrifugal pump (Bio-Console 560, Medtronic, USA), a flow sensor, an oxygenator (LivaNova, London, UK) connected to a water bath (Gentherm, USA), and a disposable TruWave pressure transducer (Edwards Lifesciences, USA) linked to the aortic cannula (Figure 1A). Organs were placed in a plastic basin (Walmart, USA) and stabilized with saline-soaked towels to prevent twisting during perfusion.

**Figure 1.**
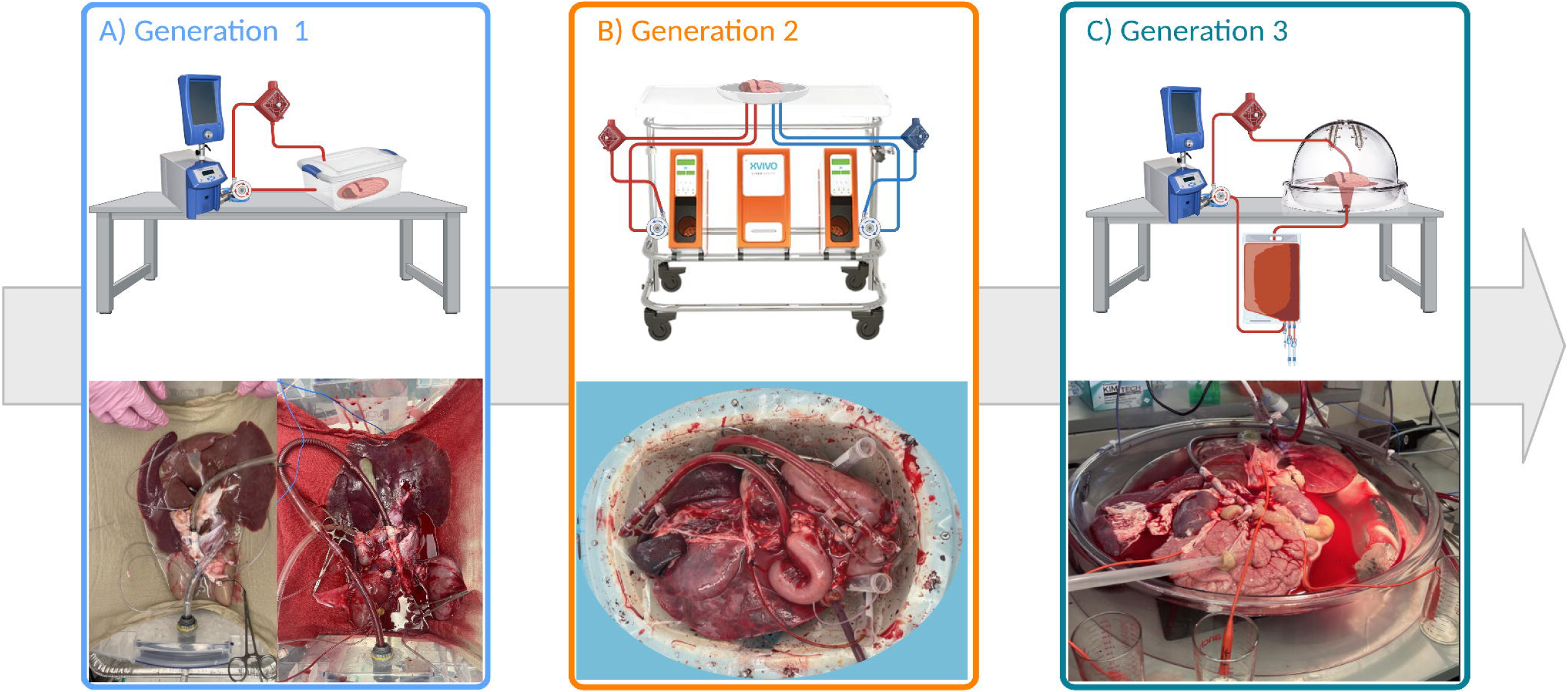
Global overview of design process and each generation of the setups used.

The multi-visceral organ blocks (n=5), containing liver, kidneys, pancreas, spleen, and a segment of the duodenum, were obtained from Yorkshire pigs at a local abattoir (following the U.S. Animal Welfare Policy), simulating a donation after circulatory death (DCD) model as extensively described in supplementary material 1. In short, blood was collected into heparinized canisters after exsanguination. A thoraco- laparotomy was performed, and the organs were flushed (supplementary material 2). The abdominal organs were removed *en bloc*, and the length of the small bowel was reduced to a 20–30 cm duodenal segment. The supra-celiac aorta was cannulated with a 28Fr cannula, and the bile duct and ureters were cannulated with 8Fr cannulas to drain bile and urine.

HMP was initiated to clear microthrombi and recharge ATP levels^17^ prior to NMP. After two hours of HMP with a glucose-enriched cold storage solution, the organs were transitioned to NMP. The perfusate for NMP consisted of autologous blood and supplements (Supplementary material 2). An infusion of sodium taurocholate (0.187 g/mL), heparin (833.33 U/mL), and Veletri (8.33 µg/mL) were all administered at 1 mL/h. Oxygenation was maintained with 100% oxygen at a flow rate of 500 mL/min.

This set-up yielded suboptimal perfusion parameters including significant perfusate loss, bowel edema, and low urine production during perfusion.

### Generation 2 Perfusion Set-Up (gen 2)

In response to the limitations observed in generation 1, the second iteration of the platform implemented several key changes. A dual perfusion system was introduced, using separate cannulas and pumps for the hepatic artery and portal vein (n=3). The Liver Assist device (XVIVO B.V., Netherlands) was repurposed for this setup, allowing for more controlled perfusion of the portal and arterial circuits (Figure 1B). Pulsitile perfusion for the aorta was introduced.

To reduce warm ischemia time and improve the quality of blood used during perfusion, organs for pig 8 were retrieved from anesthetized in-house animals rather than abattoir-sourced pigs. During procurement, the suprahepatic IVC and infra-renal aorta were cannulated to collect blood for the perfusate. The details of this procedure are extensively described in the supplementary materials (IACUC number IPROTO202300000147, Mount Sinai Hospital). Back table preparation was similar to gen 1, apart from portal vein cannulation with a 24Fr cannula.

NMP commenced after back table prep, omitting the HMP step to keep the focus on optimizing the NMP process. Dual perfusion was maintained throughout, with a separate pump controlling flow to the portal vein and aortic circuits. The perfusate composition was similar to gen 1 (Supplementary material 2).

The main limitation with this set-up was the maximum arterial flow of 1L/min afforded by the Liver Assist device. Bowel edema and fluid loss persisted, although to a lesser degree than in gen 1.

### Generation 3 Perfusion Set-Up (gen 3)

In generation 3, we reverted to single continuous perfusion through the supra-celiac aorta (n=4) (Figure 1C). The Liver Assist device was replaced with a custom-designed basin that allowed for better organ arrangement.

Organ blocks for gen 3 were similarly recovered from in-house pigs as described for gen 2. The aorta was prepared for cannulation via the thoracic aorta, and the small bowel was preserved up to 2.25 meters in length. NMP was maintained for 8–12 hours. The perfusate composition remained consistent with previous generations.

### Logging of perfusion parameters and viability assessment

Perfusate sampling was done via the arterial sample port and graft IVC, and were taken at regular intervals during perfusion while biopsies from each organ were taken at pre and post perfusion. Flow rate, mean arterial pressures (MAP), resistance, perfusate temperature, and urine output were continuously monitored and recorded. Electrolytes and blood gas values were measured via blood gas analysis using an iSTAT (ABBOTT, Chicago, United States). The oxygen consumption was calculated as followed:

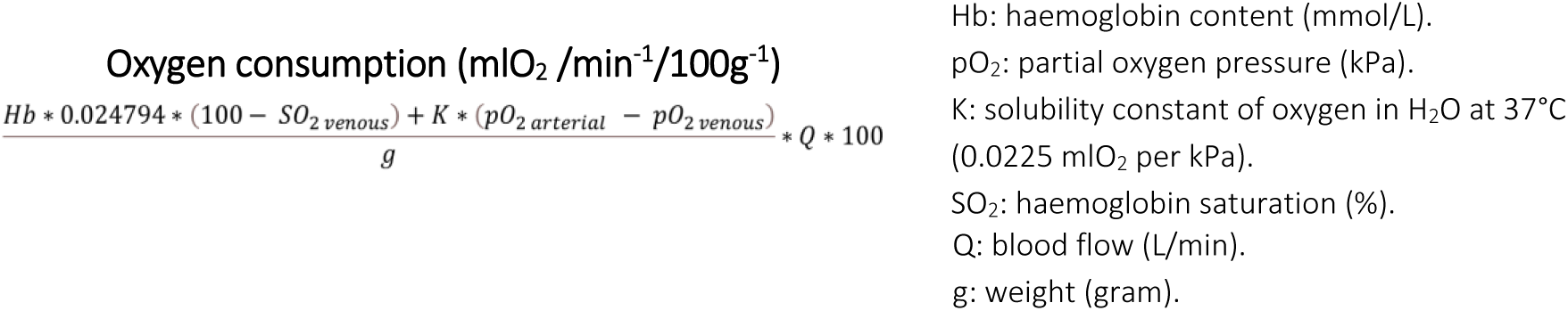

### Clinical Chemistry

Clinical chemistry analyses for alanine aminotransferase (ALT), aspartate aminotransferase (AST), alkaline phosphatase (ALP), lactate dehydrogenase (LDH), creatinine, blood urea nitrogen (BUN), potassium, lipase, and amylase were conducted routinely.

### Histological analysis

Biopsies were obtained from the renal cortex, liver parenchyma, pancreas, and the luminal wall of the duodenum, and fixed in 4% formalin. These samples were embedded in paraffin wax, sectioned at 4 µm, and stained with hematoxylin and eosin (H&E) to highlight morphological features. Histological preparation was performed, sections were evaluated and scored in a blinded manner by a board-certified pathologist using established scoring protocols.^18–22^

### Data and statistical analysis

GraphPad Prism (version 9.1.0) was used to visualize the data and perform statistical analysis. Longitudinal perfusion data per generation are shown as single values over time in the figures and the overall arithmetic mean with the standard deviation (SD) as described in Table 2. Differences across experimental groups were analyzed using one-way analysis of variance (ANOVA) followed by Tukey’s multiple comparisons. Injury markers are expressed as aligned scatter plots and the arithmetic mean with the SD. Differences across experimental groups were assessed using a two-way ANOVA with Fishers LSD test. All statistical tests were two-tailed, and differences between groups were considered statistically significant when *p* < 0.05.

**Table 1.** Preservation characteristics for each group shown as the mean+SD.

| Demographics | Experimental groups |  |  | P value |  |  |
| --- | --- | --- | --- | --- | --- | --- |
|  | Gen 1 | Gen 2 | Gen 3 | G1 vs. G2 | G1 vs. G3 | G2 vs. G3 |
| **WIT (min)** | 9.4 ± 3.1 | 8.3 ± 1.2 | 3.3 ± 1.7 | 0.8086 | 0.0085 | 0.0447 |
| **Cold flush (min)** | 11.8 ± 0.8 | 18.7 ± 13.9 | 17.3 ± 2.5 | 0.3812 | 0.4774 | 0.9590 |
| **CIT (min)** | 368.6 ± 55.8 | 265.3 ± 61.9 | 184.8 ± 32.3 | 0.0506 | 0.0011 | 0.1497 |
| **Duration of HMP (min)** | 110.8 ± 21.78 | - | - | - | - | - |
| **Duration of NMP (min)** | 438.0 ± 47.6 | 440.0 ± 69.3 | 480.0 ± 176.6 | 0.9997 | 0.8438 | 0.8873 |
| **Mean MAP during NMP (mmHg)** | 110.3 ± 22.31 | 73.15 ± 13.32 | 63.60 ± 7.565 | <0.0001 | <0.0001 | 0.0956 |

WIT: warm ischemia time, CIT: cold ischemia time, HMP: hypothermic machine perfusion, NMP: normothermic machine perfusion, MAP: mean arterial pressure.

**Table 2.** Mean aortic flow, resistance, total urine output, oxygen consumption, lactate clearance and perfusate pH of grafts in the different experimental groups over time. Shown as mean+SD.

| NMP parameters | Experimental groups |  |  | P value |  |  |
| --- | --- | --- | --- | --- | --- | --- |
|  | *Gen 1* | *Gen 2* | *Gen 3* | *G1 vG. G2* | *G1 vG. G3* | *G2 vG. G3* |
| Aortic flow (L/min) | 1.018 ± 0.506 | 0.617 ± 0.201 | 2.089 ± 0.633 | 0.0498 | <0.0001 | <0.0001 |
| Aortic resistance | 0.224 ± 0.127 | 0.195 ± 0.148 | 0.038 ± 0.027 | 0.6916 | <0.0001 | <0.0001 |
| Total urine output (mL) | 51.90 ± 99.97 | 2 ± 3.46 | 271.3 ± 324.8 | 0.9375 | 0.278 | 0.233 |
| Oxygen consumption (mL O<sub>2</sub>/min) | 43.56 ± 16.57 | 19.42 ± 5.21 | 49.52 ± 15.79 | 0.0082 | 0.6583 | 0.0009 |
| Lactate clearance (mmol/L) | 10.44 ± 3.72 | 3.26 ± 2.37 | 3.10 ± 2.21 | <0.0001 | <0.0001 | 0.993 |
| Perfusate pH | 7.27 ± 0.34 | 7.29 ± 0.14 | 7.30 ± 0.13 | 0.9843 | 0.961 | 0.9961 |

## Results

### Preservation characteristics

The preservation parameters for each pig are summarized in Table 1. Gen 3 achieved a significant reduction in warm ischemia time (WIT) compared to gens 1 and 2. Cold ischemia time was notably longer in gen 1 than in gens 2 and 3. HMP was performed exclusively in gen 1. No significant differences were observed in the durations of NMP across the different generations. However, the MAP during NMP in gen 1 was significantly higher compared to the other generations.

### Generation 1 perfusion

In the initial experimental setup, oxygenated HMP was first performed followed by NMP. The perfusion parameters for HMP are detailed in Figures 2A-C. Aortic resistance steadily decreased, reaching 0.028 ± 0.006 by the end of the 120-minute HMP period (Figure 2B). Hemodynamic parameters during NMP are presented in Figures 2D-E. The grafts exhibited physiological flow rates, with a mean of 1.018 ± 0.506 L/min (Figure 2D). Except for pigs 1 and 5, resistance generally decreased over time (Figure 2E). Urine production was generally low, except for pig 1, with an average cumulative output of 51.90 ± 99.97 mL (Figure 2F). Oxygen consumption fluctuated throughout the NMP period (Figure 2G), while perfusate lactate levels increased (Figure 2H) and perfusate pH progressively declined (Figure 2I), indicating suboptimal perfusion.

**Figure 2.**
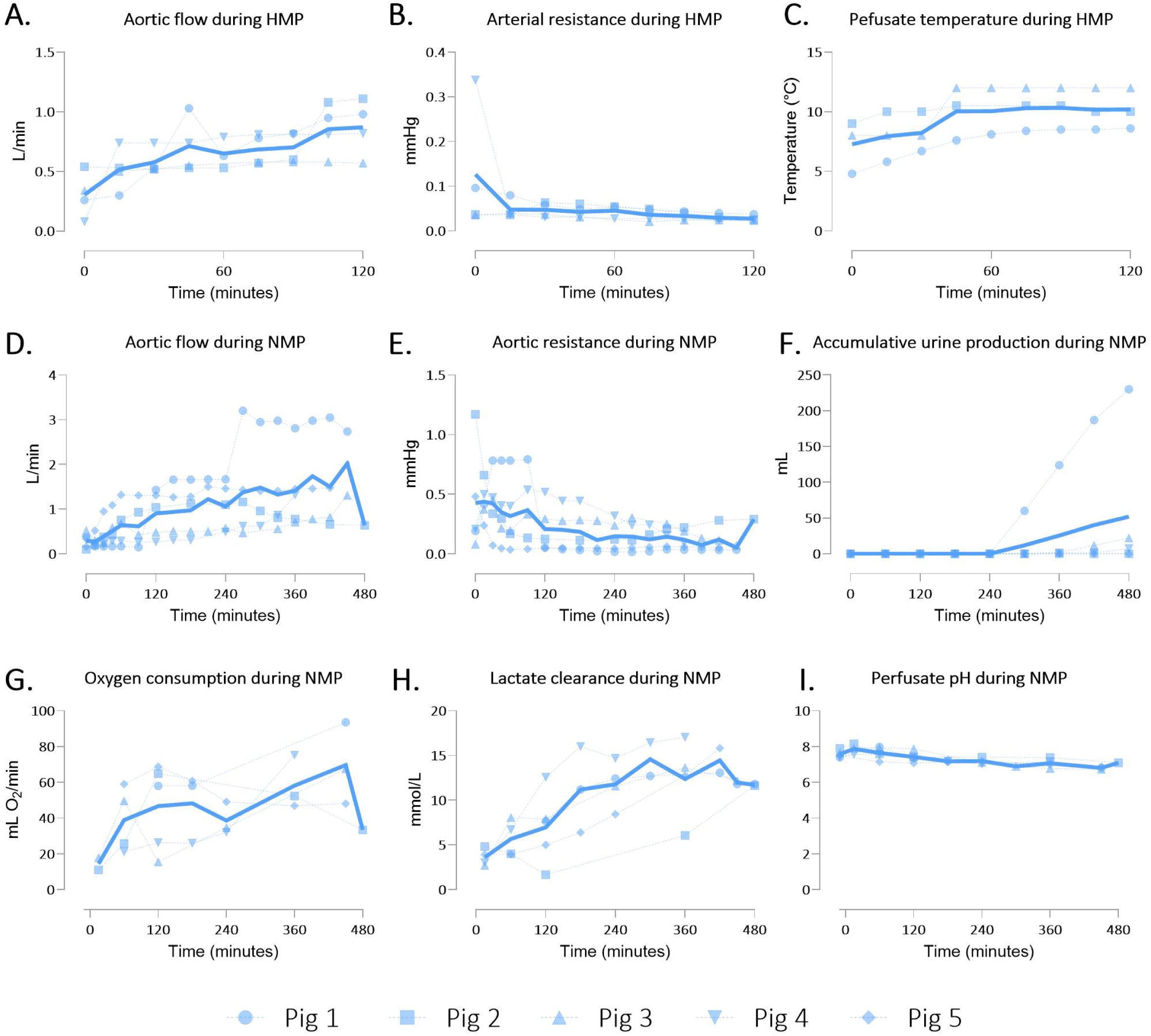
Perfusion parameters of multivisceral grafts in gen 1. A) Aortic flow, B) aortic resistance, and C) graft temperature during hypothermic machine perfusion (HMP). D) Aortic flow, E) aortic resistance, F) accumulative urine production G) oxygen consumption H) lactate clearance and I) perfusate pH during normothermic machine perfusion (NMP). Data shown as individual values (dotted lines) and mean (continuous line).

### Generation 2 perfusion

Given the rising lactate levels observed in the initial generation, we hypothesized that perfusion was inadequate. The shortening of the bowel may have reduced mesenteric venous return leading to portal hypo-perfusion. To address these issues, NMP was performed with dual arterial and portal perfusion. In this generation, the mean aortic flow during perfusion was 0.617 ± 0.201 L/min, constrained to a maximum of 1 L/min due to machine limitations, resulting in significantly lower flow rates compared to the first gen (Figure 3A/Table 2). The mean portal flow was 0.427 ± 0.228 L/min (Figure 3A). Aortic and portal resistances were 0.195 ± 0.148 and 0.062 ± 0.170, respectively, throughout the perfusion (Figure 3B). Notably, only Pig 6 produced urine (Figure 3C). Oxygen consumption fluctuated over time but was significantly lower than in gen 1 (Figure 3D/Table 2). Lactate clearance was achieved in two of the three pigs, with mean lactate levels significantly lower compared to gen 1 (Figure 3E/Table 2). The perfusate pH remained stable throughout the perfusion (Figure 3F). Additionally, bowel peristalsis was observed in the shortened segments during all three perfusions (Supplementary video 1).

**Figure 3.**
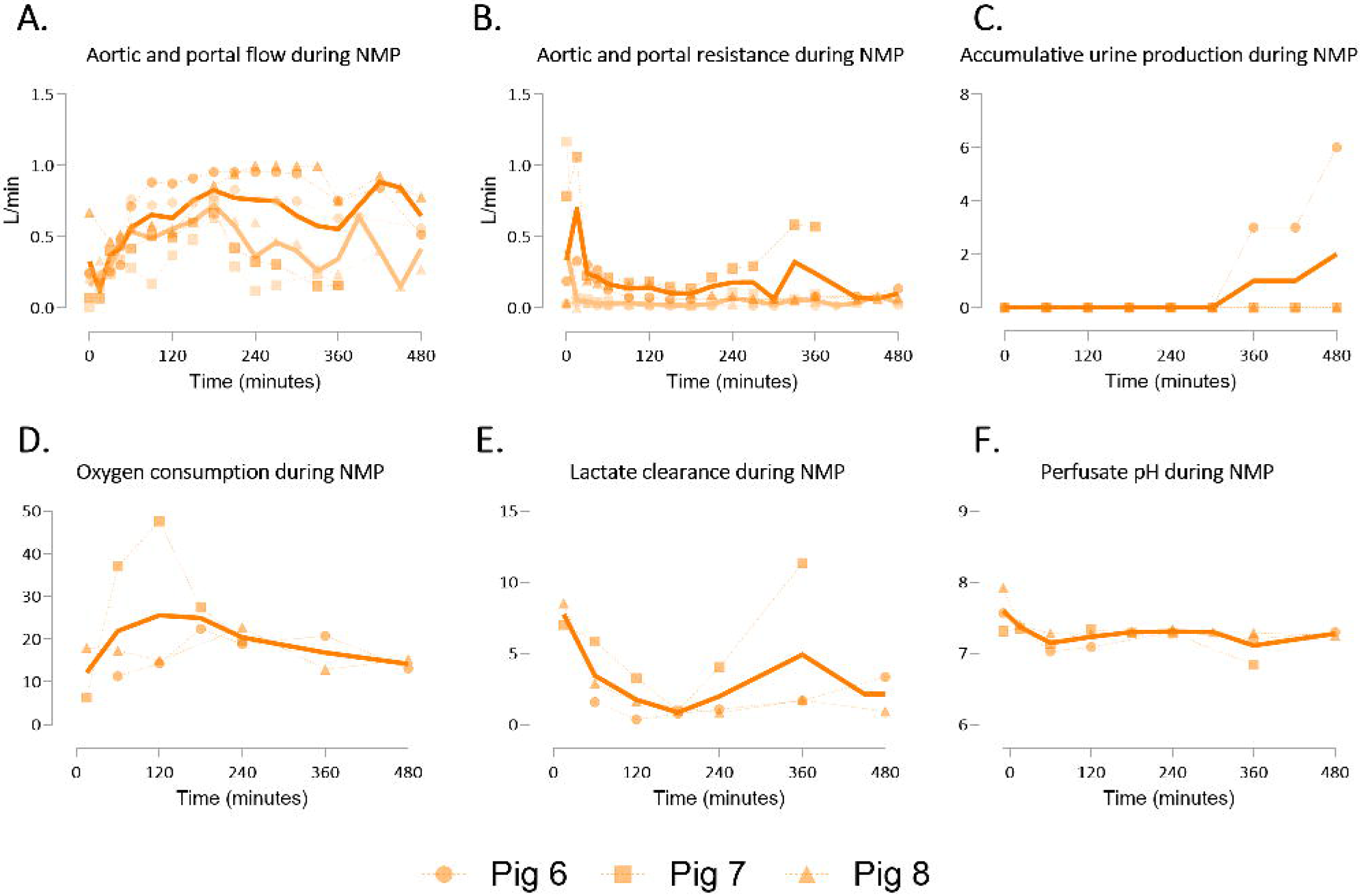
Perfusion parameters of multivisceral grafts in gen 2. A) • Aortic flow and • portal flow, B) • aortic and • portal resistance, and C) accumulative urine production D) oxygen consumption E) lactate clearance and F) perfusate pH during normothermic machine perfusion (NMP). Data shown as individual values. Data shown as individual values (dotted lines) and mean (continuous line).

### Generation 3 perfusion

In the third experimental generation, single continuous perfusion was utilized while maintaining a longer section of the small bowel intact (2.25m) to reduce the risk of portal hypoperfusion. The mean aortic flow during perfusion was 2.089 ± 0.633 L/min, significantly higher than in gens 1 and 2 (Figure 4A/Table 2). Aortic resistance throughout perfusion was 0.038 ± 0.027, markedly lower compared to the previous gens (Figure 4B/Table 2). Urine production was observed in all pigs (Figure 4C). Oxygen consumption fluctuated over time and was significantly higher than in gen 2 (Figure 4D/Table 2). Lactate clearance was achieved in four of the five pigs, with mean lactate levels significantly lower than in gen 1 (Figure 4E/Table 2). The perfusate pH remained stable throughout the perfusion period (Figure 4F). Additionally, bowel peristalsis was observed during all perfusions (Supplementary video 2).

**Figure 4.**
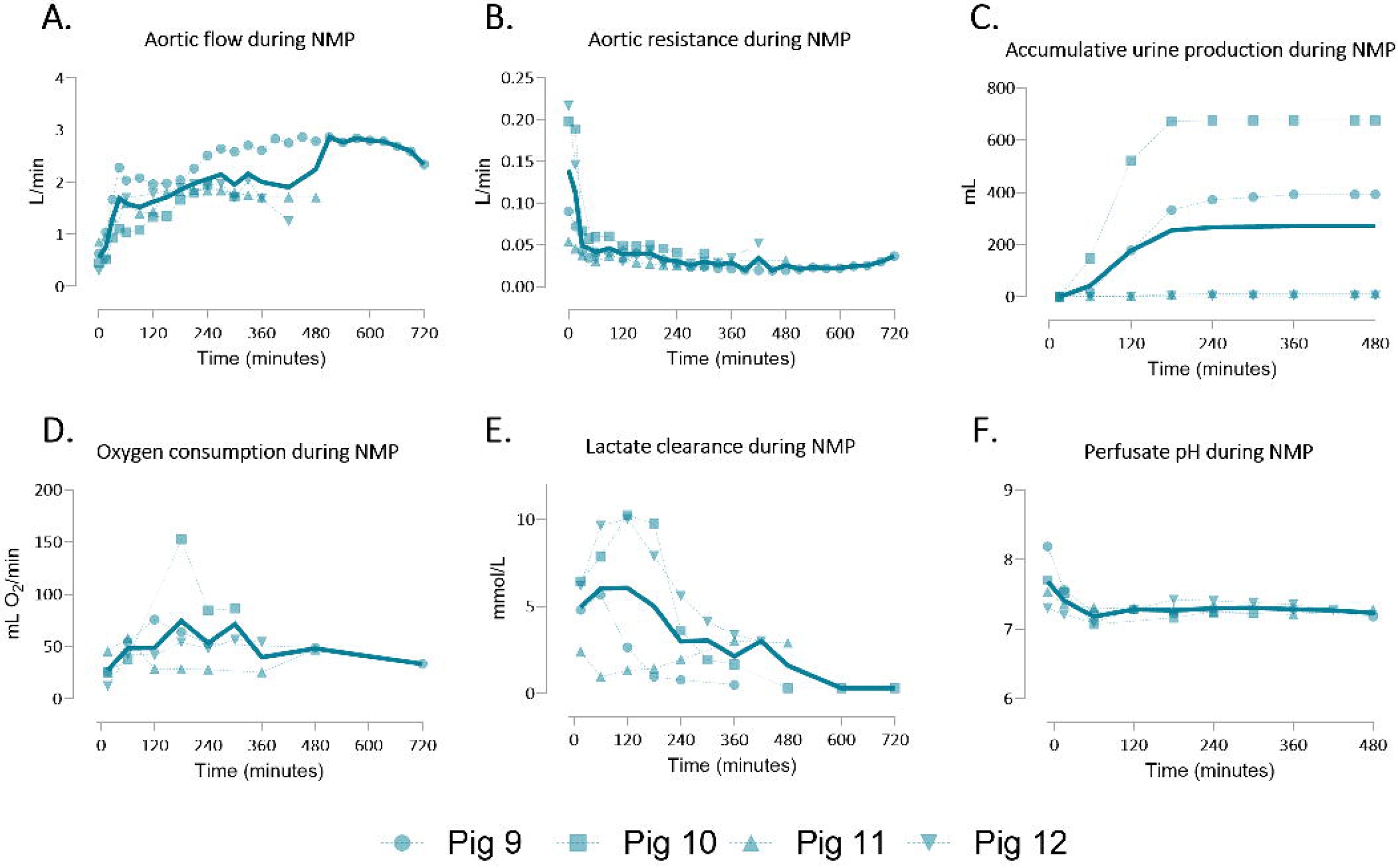
Perfusion parameters of multivisceral grafts using gen 3. A) Aortic flow, B) aortic resistance, and C) accumulative urine production D) oxygen consumption E) lactate clearance and F) perfusate pH during normothermic machine perfusion (NMP). Data shown as individual values.

### Cellular injury during multivisceral perfusion

To assess cellular injury during multi-visceral perfusion, we evaluated a comprehensive panel of biomarkers. Liver function was analyzed by measuring ALT, AST, and ALP levels (Figures 5A-C). AST levels significantly increased in gen 1 and 3 (Figure 5B). Although not statistically significant, the pigs in gen 3 exhibited the lowest ALT and AST levels at the end of perfusion. ALP levels remained relatively stable, except in pig 7, where an increase was observed (Figure 5C). LDH, a marker of tissue damage, also significantly increased during perfusion in gen 1 and 3 (Figure 5D). However, there were significantly lower levels at the end of perfusion in gen 3 compared to gen 1. To evaluate renal function, creatinine was added to the perfusate to measure creatinine clearance. Perfusate creatinine levels significantly decreased during perfusion in gen 3(Figure 5E). Blood urea nitrogen (BUN) levels, a marker of glomerular filtration, showed a significant increase over time in gen 3 (Figure 5F). Perfusate potassium levels significantly decreased over time in gen 3, and were significantly lower compared to gen 1 (Figure 5G). Additionally, lipase and amylase were measured as markers of pancreatic function (Figures 5H-I). Amylase was significantly higher after 240 minutes of perfusion in gen 3, but not at the end of perfusion.

**Figure 5.**
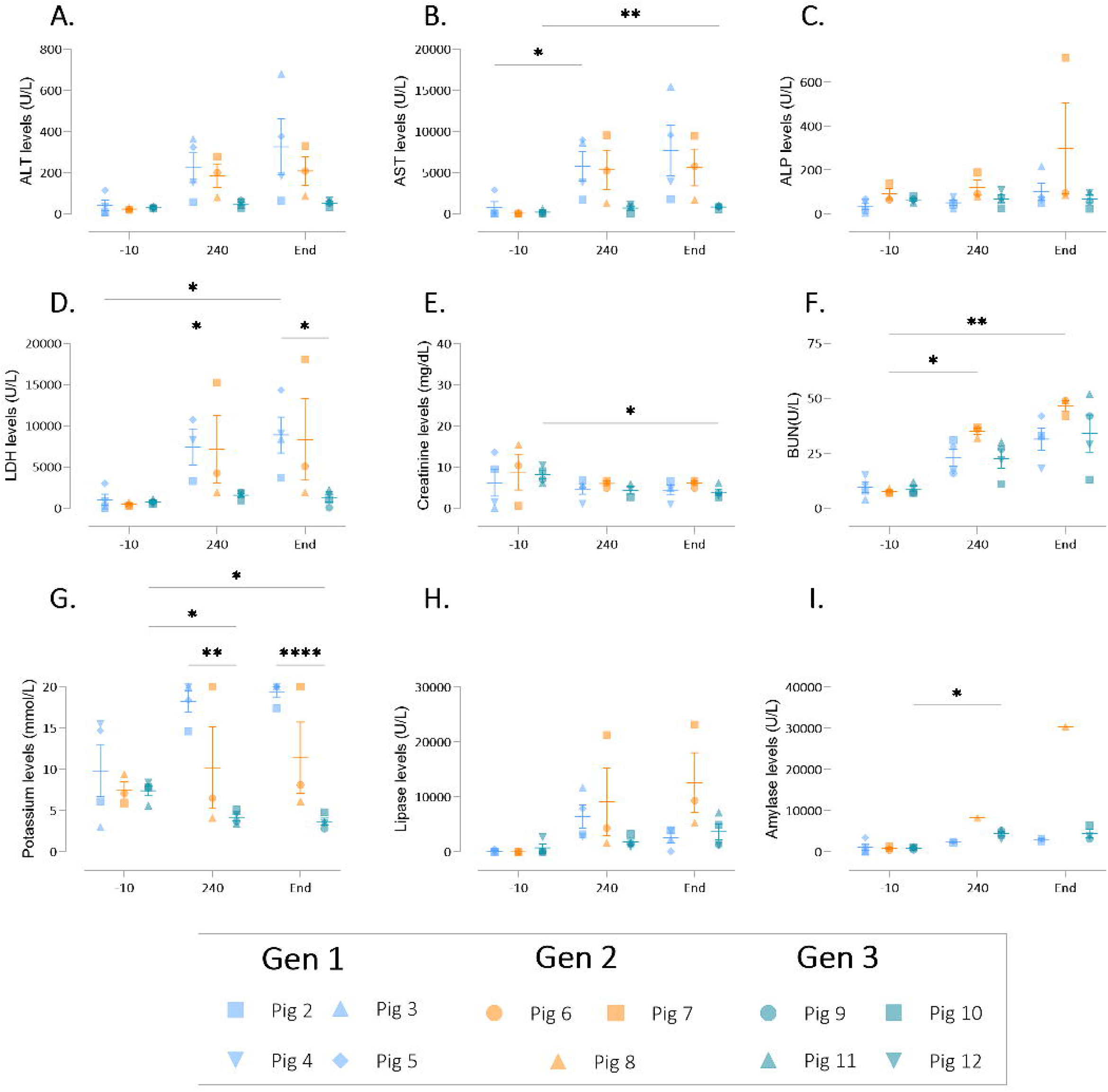
Cellular injury during multi-visceral perfusion. A) alanine aminotransferase (ALT), B) aspartate aminotransferase (AST), C) and alkaline phosphatase (ALP), D) Lactate dehydrogenase (LDH), E) creatinine, F) blood urea nitrogen (BUN), G) potassium, H) Lipase and I) Amylase were measured at baseline (-10), 4 hours into perfusion (240), and at the end of perfusion (end). Data are expressed as aligned scatter plots and the arithmetic mean+SD. * *p* < 0.05, ** *p* < 0.01.

### Histological changes over time

Histological changes in the liver, kidneys, pancreas, and intestine were analyzed both before and after perfusion (Figure 6-8). Notable differences in liver tissue were primarily observed in gen 1, in which histological scoring using the Modified Hansen Score^18^ revealed significantly greater biliary mucosal loss, hepatic arteriosclerosis, and mural stroma necrosis following perfusion (Figure 6B/C/E). Additionally, gen 1 resulted in increased and hepatic necrosis, although these increases were not statistically significant (Figure 6D/G). In gens 2 and 3, a non-significant increase in biliary inflammation and hepatic necrosis was also noted (Figure 6D/G).

**Figure 6.**
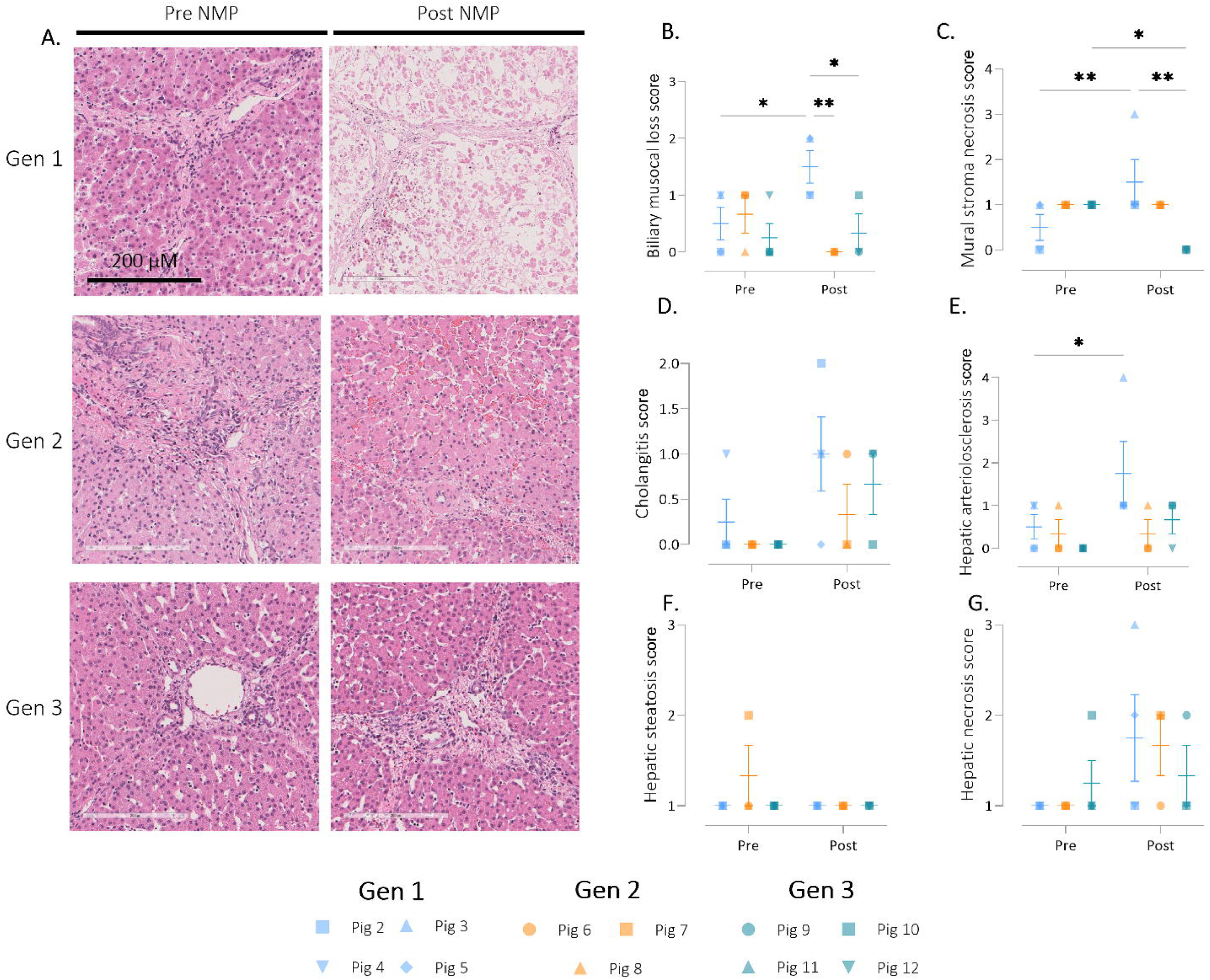
Histological assessment of the liver during multi-visceral perfusion. A) Liver parenchyma hematoxylin and eosin-stained sections representing pre and post NMP (Gen 1 = pig 3, Gen 2 = pig 8 and Gen 3 = pig 12). B) biliary mucosal loss, C) mural stromal necrosis, D) cholangitis, E) hepatic arteriosclerosis, F) hepatic steatosis G) hepatic necrosis were scored. Data are expressed as aligned scatter plots and the arithmetic mean+SD. * *p* < 0.05, ** *p* < 0.01. Original magnification: 20X, scale bar is 200µM.

**Figure 7.**
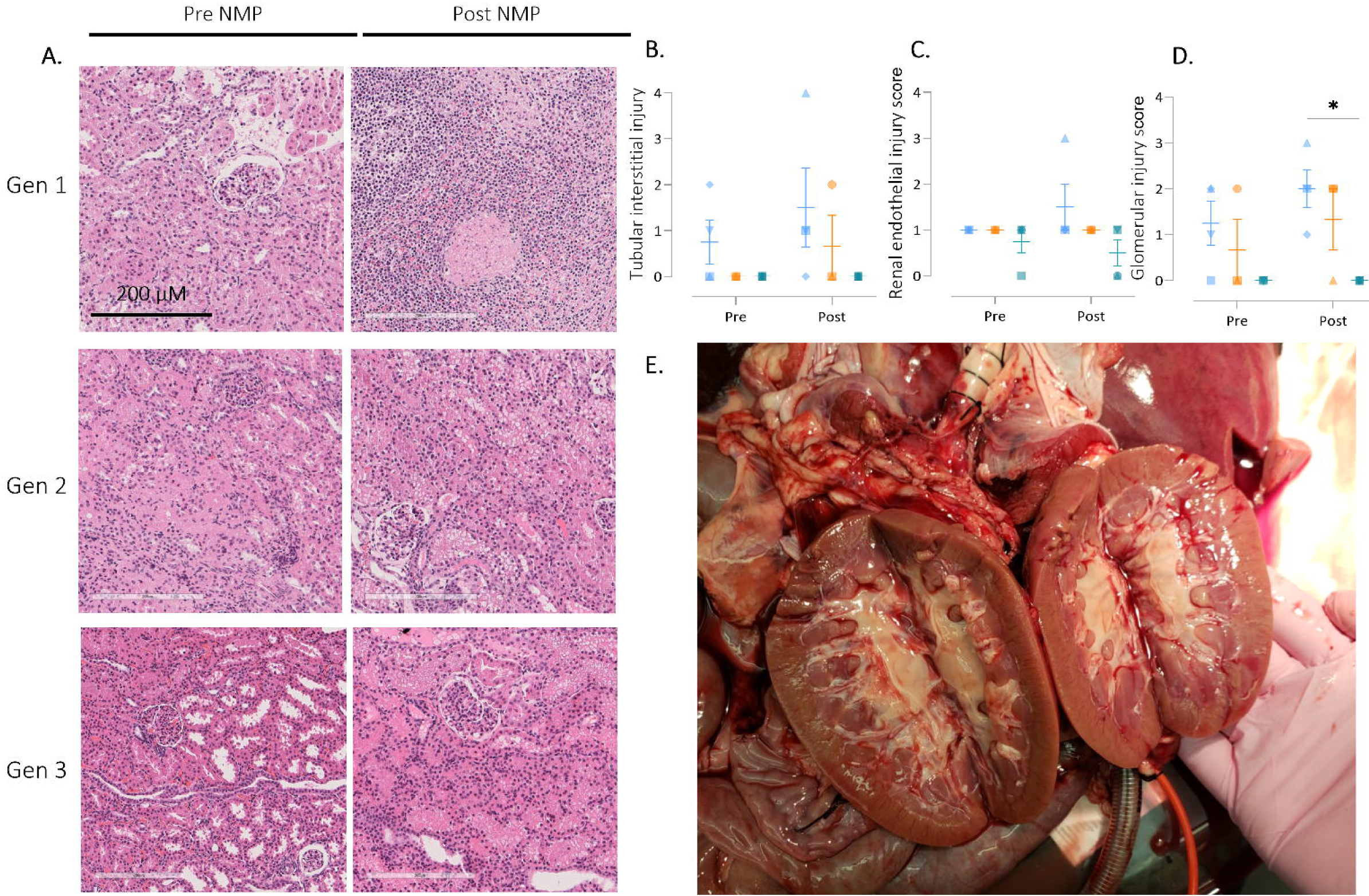
Histological assessment of the kidney during multi-visceral perfusion using the EGTI scoring system. A)Renal cortex hematoxylin and eosin-stained sections representing pre and post NMP (Gen 1 = pig 3, Gen 2 = pig 8 and Gen 3 = pig 12). B) glomerular injury score C) renal endothelial injury D) tubular interstitial were scored. E) macroscopic image of the kidneys post NMP (pig 12). Data are expressed as aligned scatter plots and the arithmetic mean+SD. Original magnification: 20X, scale bar is 200µM.

**Figure 8.**
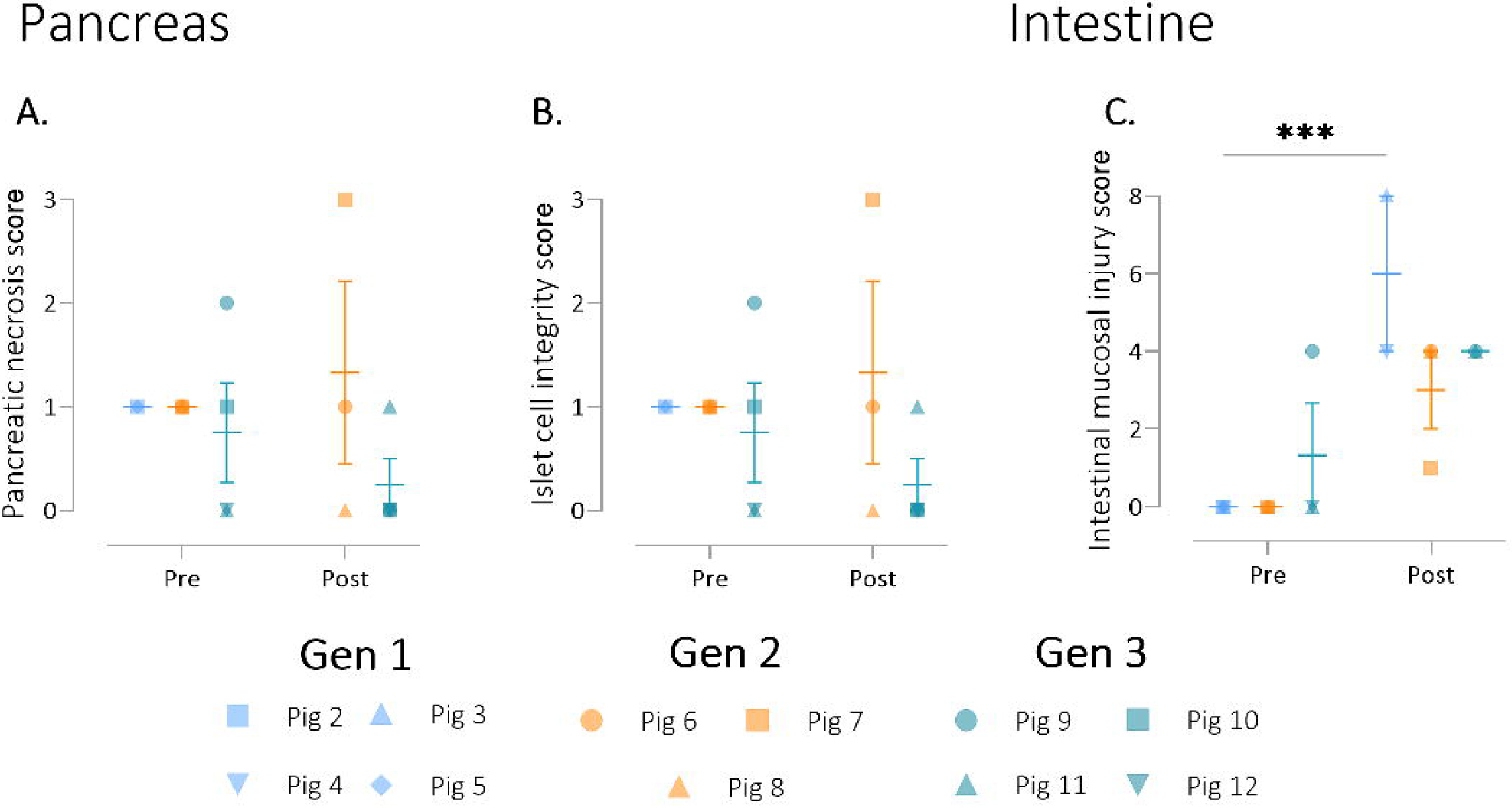
Histological assessment of the Pancreas and intestine during multi-visceral perfusion. A) pancreatic necrosis score, B) Islet cell integrity, and C) Intestinal musical injury were scored pre and post NMP. Data are expressed as aligned scatter plots and the arithmetic mean+SD. ** *p* < 0.01.

Histological evaluation of the kidneys using the EGTI scoring system^21^ displayed more inflammation in gen 1 and minimal edema in all groups post perfusion (Figure 7A). No differences were observed when looking at tubulo-interstitial and endothelial injury (Figure 7B/C). A significantly higher glomerular injury score in gen 1 compared to gen 3 was observed (Figure 7D). Macroscopically, the kidneys looked pink and felt soft post NMP (Figure 7E).

No significant changes were observed in the pancreas, although fat/parenchymal necrosis and islet cell integrity appeared to improve following perfusion in gen 3 (Figure 8A/B). Lastly, intestinal mucosal injury, as assessed by the Chiu/Park score,^22^ significantly worsened after perfusion in gen 1 (Figure 8C).

## Discussion

The organ shortage highlights the urgent need for advancements in machine preservation techniques, particularly in normothermic perfusion of kidney, pancreas, and small bowel. This study aimed to develop a multivisceral perfusion platform that systematically characterizes perfusion dynamics, physiological parameters, and molecular and histological outcomes through an iterative design process. We demonstrated the feasibility of abdominal multivisceral NMP for up to 8 hours, achieving stable arterial flow, perfusate pH, and sufficient oxygen consumption, all indicating organ viability.

A key strength of our study lies in the iterative design approach, which allowed for continuous, data-driven refinements. Unlike a traditional linear design process, in which designs are finalized before testing, our approach enabled real-time problem-solving. For instance, after identifying excessive bowel edema and fluid loss in gen 1, we introduced significant changes in gen 2, including shortening the small bowel and implementing dual-circuit perfusion for the hepatic and portal systems. These adjustments improved portal flow, and enhanced lactate clearance. Gen 3 further improved outcomes by adopting single continuous perfusion with higher flow rates and a redesigned basin for optimal organ arrangement. This process not only yielded better results but also reduced the number of animals required by addressing issues early on.

The iterative design of our system provided the flexibility to optimize critical parameters, including flow rates, pressures, and organ stability, facilitating the standardization of multivisceral perfusion protocols.

The optimal perfusion parameters for multivisceral abdominal perfusion remain undefined. However, we identified specific conditions that support multi-organ viability during NMP. In our third-generation system, an arterial flow rate of 0.8-1.7 L/min resulted in sufficient perfusion, as evidenced by effective lactate clearance and stable perfusate pH. Notably, single arterial perfusion exhibited superior perfusion characteristics compared to dual arterial and portal perfusion. The latter was potentially limited by the 1 L/min arterial flow used in this setup. The ideal MAP during NMP also remains unclear, as it is uncertain whether all abdominal organs benefit from uniform arterial pressure. In our study, maintaining the MAP between 60-75 mmHg supported adequate tissue oxygenation, urine production, and biochemical stability. It is likely that certain organs, such as the pancreas or small intestine, may require individualized arterial pressure modulation to optimize viability during NMP. Further research is warranted to investigate this aspect.

While urine production improved in gen 3, the composition of urine and its relation to kidney histology were not fully evaluated. We did observe a decrease in perfusate creatinine levels, while BUN levels only increased slightly, reflecting proper kidney function. Furthermore, the post NMP kidney histology and macroscopic appearance remained similar to pre NMP.

Amylase and lipase levels rose slightly, with no histological signs of pancreatic injury post-NMP, suggesting minimal pancreatic damage. These levels were considerably lower than those reported in isolated porcine pancreas perfusion.^23–25^ Currently, NMP of pancreas grafts has been minimally explored, with perfusion times often restricted by graft edema and injury.^10,20^ Pancreas perfusion remains underexplored, but our results indicate that multivisceral abdominal perfusion may offer an effective approach for preserving the pancreas *ex vivo*.

Additionally, intestinal peristalsis was consistently observed in gen 3, a key indicator of intestinal function.^26^ Peristalsis, the rhythmic contractions of the gut wall, is intrinsic to the gastrointestinal tract and persists even without external autonomic control. While it has been documented in hypothermic and sub- normothermic isolated intestine perfusion, its occurrence during normothermic perfusion has not been widely reported.^12,27^ Histological analysis revealed increased intestinal injury scores post-NMP, although previous studies on isolated porcine intestinal perfusion showed lower injury scores with shorter (6-hour) perfusions at sub-normothermic temperatures.^11,12^ The observed bowel edema and significant fluid loss in our study limited perfusion to 7-12 hours. We hypothesize that this fluid loss stems from compromised luminal integrity caused by ischemia-reperfusion injury (IRI). Intestinal grafts from donors after circulatory death (DCD) are particularly susceptible to IRI, making them generally non-viable for transplantation.^28^ Since all grafts in this study were exposed to WIT, this likely contributed to the intestinal injury. Potential improvements for intestinal preservation include a cold luminal flush, extended bowel length, better flow and pressure control, and nutrient supplementation during perfusion.

The feasibility of multivisceral perfusion was first demonstrated in studies by Prieto et al. (1988)^13^ and Chien et al. (1989)^14^, in which stable perfusion of multiple organs (heart, lungs, liver, kidneys, pancreas) was achieved for up to 22 and 37 hours, respectively. More recent studies by Chen et al. (2021)^15^ and Stevens et al. (2024)^16^ have explored perfusing mini-pig and butcher-pig abdominal organs for 7-10 hours using dual-circuit perfusion. However, these studies involved significant variations in perfusion protocols, with the early studies focusing on heart and lung perfusion at lower temperatures and minimal intestinal involvement. In contrast, Chien did not subject grafts to cold ischemia, while Stevens worked with butcher pigs, during which significant warm ischemia is inevitable. Moreover, none of these studies reported detailed histological assessments, which are critical for evaluating tissue integrity. Both Chien and Stevens perfused longer bowel segments and observed bowel edema along with intestinal fluid loss. Moreover, none of the groups demonstrated lactate clearance, and all reported elevated ALT and AST levels, indicating suboptimal liver viability during multivisceral perfusion.

Despite the valuable insights gained from this study, several limitations should be acknowledged. Early experiments using abattoir-sourced pigs introduced variability in ischemia times and perfusate quality, complicating comparisons across experimental generations. The decision to use abattoir pigs aimed to minimize animal suffering during the initial exploratory phase. Early biopsies of the pancreas and intestine were of lesser quality. Additionally, the small sample size reflects the significant resources required for these experiments, and limited blood volume restricted perfusion duration due to fluid losses.

Our third-generation platform achieved superior outcomes, with improved perfusion parameters, reduced tissue damage, and enhanced organ function. This suggests that a single continuous perfusion strategy, incorporating an intact bowel segment, may be the most effective approach for maintaining organ viability during multivisceral perfusion. Further research is necessary to optimize this technique, but multivisceral perfusion holds significant potential for improving organ preservation and enabling resuscitation prior to transplantation. Extending perfusion times to 2-3 weeks could potentially allow for "organs on demand," addressing the pressing time constraints of organ allocation and helping bridge the gap between supply and demand in transplantation, ultimately improving patient outcomes.

## Acknowledgements/Funding

This work was generously supported by the Blavatnik family foundation. We would like to thank Jamshid Abdul-ghafar for his expert histological assessment and Avery Fortier, Kimberly Feeney, Sophia Gamboa, Cole Brown, Brittany Sacks, Quinn Bogue, Andrew Rosowicz, Ahmed Hassan, and Jackson Breaks for their assistance with the experiments and insightful comments.

## Data availability statement

Data available upon request.

## IACUC statement

All procedures involving laboratory animals were reviewed and approved by the Institutional Animal Care and Use Committee of the Icahn School of Medicine at Mount Sinai (IACUC protocol number: IPROTO202300000147).

## Supporting information statement

Additional supporting information may be found online in the Supporting Information section”

## Abbreviation page

ALP: Alkaline phosphatase
ALT: Alanine aminotransferase
ANOVA: Analysis of variance
AST: Aspartate aminotransferase
BUN: Blood urea nitrogen
DCD: Donation after circulatory death
H&E: Hematoxylin and eosin
HMP: Hypothermic machine perfusion
IRI: Ischemia-reperfusion injury
LDH: Lactate dehydrogenase
MAP: Mean arterial pressure
NMP: Normothermic machine perfusion
WIT: Warm ischemia time

