## Supplementary material for "An iterative design approach to development of an ex-vivo normothermic multivisceral perfusion platform"

### Supplementary material 1

#### *Organ procurement at the abattoir*

Multivisceral abdominal organ blocks containing liver, kidneys, pancreas, spleen and segment of the duodenum were recovered from Yorkshire pigs from a local abattoir following the U.S. Animal Welfare Policy. Following exsanguination, blood was collected in heparinized (25,000 IU) canisters. Afterwards, the pig was positioned supine on the operative table and rapidly prepared for surgical procurement. A thoraco-laparotomy was performed, followed by swift cannulation of the aorta and the initiation of cold perfusion in the abdomen. The abdominal inferior vena cava (IVC) was vented, and a cross-clamp was applied to the descending thoracic aorta. The organs were flushed with ice-cold flush solution (supplementary material 2), and the abdominal and thoracic cavities were packed with ice for topical cooling. The flush was continued until the flush returned clear, signaling the cessation of perfusion.

#### *Back-table preparation*

Evisceration of the organs was then performed. The rectum was transected, and the ureters were identified and marked. The infra-renal IVC and aorta were transected in the abdomen. Cranial dissection was carried out via the retroperitoneum, allowing the entire intestine, both kidneys, spleen, liver, pancreas, and associated bowel mesentery to be taken *en bloc*. The thoracic aorta was transected below the level of the cross-clamp leaving sufficient length on the thoracic aorta for cannulation, and the suprahepatic IVC was divided. The esophagus was transected and organ bloc explanted.

After explantation, the large bowel was resected, while the small bowel was left intact but shortened to minimize edema and insensible losses during perfusion. The stomach was resected at the level of the pylorus leaving a short remnant gastric pouch. The aorta was prepared for cannulation via the thoracic aorta, with a 28Fr cannula placed (Figure 1). Lumbar arteries were identified and ligated. A leak test was performed to ensure no arterial leaks from the aorta or its branches before perfusion. The ureters and common bile duct were cannulated with 8Fr cannulas (Figure 1), and the cystic duct was identified and ligated. The distal end of the bowel was cannulated with a 30 Fr catheter to allow for the escape of effluent (Figure 1).

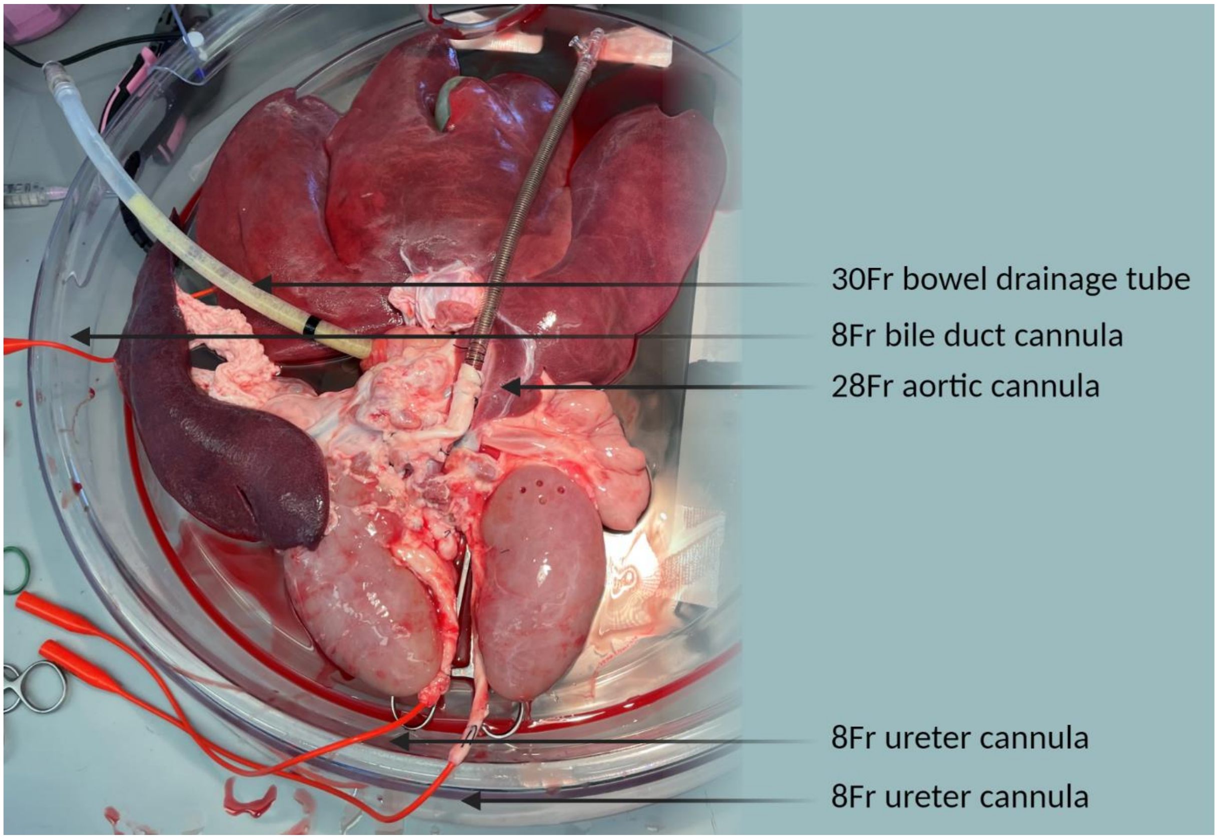


**Figure 1**. cannulation during multivisceral perfusion.

*Surgical protocol and organ procurement at the in-house animal facility*

The preparation and surgical procedure for the animals prior to the experimental procedures was similar for all pigs. The animals were fasted for 24 hours prior to surgery. The animals were sedated with tiletamine-zolazepam at a dosage of 5-6 mg/kg. An intravenous (IV) catheter was then placed via the ear vein, and vital signs, including heart rate, $SpO_2$, and temperature, were collected. Based on the animal's size and weight, an appropriately sized endotracheal tube is selected. The animal was then intubated in the supine position, and the tube is secured to ensure a patent airway. After intubation, the animal was transferred to the operating room, and connected to sensors to continuously monitor heart rate, $SpO_2$, and respiration. Finally, the animal was placed on ventilation with isoflurane at a concentration of 1-3% and oxygen at 2.5 L/min. All animals were euthanized while under general anesthesia.

A thoraco-laparotomy was performed, and 30,000 units of heparin were administered to ensure complete heparinization before blood collection. The infra-renal aorta was encircled, and the supra-hepatic cava was prepared for cannulation. The blood was collected in a controlled manner by passive drainage into a sterile collection bag pre-lined with 25,000 units of heparin, aided by tilt augmentation of the pig. After confirming death, a flush was initiated (see supplementary material 2). The remainder of the procurement process was consistent with previous descriptions.

Supplementary material 2

| | Pig 1 | Pig 2 | Pig 3 | Pig 4 | Pig 5 | Pig 6 | Pig 7 | Pig 8 | Pig 9 | Pig 10 | Pig 11 | Pig 12 |
|---|---|---|---|---|---|---|---|---|---|---|---|---|
| Generation set-up | 1 | 1 | 1 | 1 | 1 | 2 | 2 | 2 | 3 | 3 | 3 | 3 |
| Pig type | Butcher | Butcher | Butcher | Butcher | Butcher | Butcher | Butcher | Laboratory | Laboratory | Laboratory | Laboratory | Laboratory |
| Pig age (months) | 6 | 6 | 8 | 6 | 6 | 6 | 5 | 3 | 3.5 | 3 | 3 | 3 |
| Pig sex | Female | Female | Female | Male | Female | Male | Male | Male | Male | Male | Male | Male |
| Pig weight (Kg) | NR | NR | NR | NR | NR | NR | NR | 46.4 | 50.9 | 33.8 | 37 | 39.9 |
| Flush type | 3L NaCl + 4L HTK | 3L NaCl + 4L Servator B™ Safe | 3L NaCl + 4L HTK | 3L NaCl + 4L HTK | 3L NaCl + 4L HTK | 6L NaCl | 3L NaCl + 4L HTK | 3L NaCl + 4L HTK | 3L NaCl + 4L HTK | 3L NaCl + 4L HTK | 6L HTK | 6L HTK |
| CIT (hours) | 5:55 | 5:45 | 5:57 | 5:21 | 7:45 | 4:58 | 5:04 | 3:14 | 2:52 | 3:29 | 3:33 | 2:25 |
| Bowel length (m) | 0.03 | 0.03 | 0.02 | 0.1-0.3 | 0.1-0.3 | 0.03 | 0.03 | 0.03 | 2.25 | 2.25 | 2.25 | 2.25 |
| Graft weight pre NMP (g) | NR | 1013 | 1453 | NR | NR | NR | NR | 1900 | 1800 | 2550 | 1800 | 2000 |
| *Perfusate composition NMP* | | | | | | | | | | | | |
| Whole blood (mL) | 1000 | 1000 | 1000 | 1600 | 1900 | 1250 | 1300 | 1800 | 1775 | 2100 | 1550 | 1670 |
| Cefuroxim in 20 mL naCl (g) | 1.5 | 1.5 | 1.5 | 1.5 | 1.5 | 1.5 | 1.5 | 1.5 | 1.5 | 1.5 | 1.5 | 1.5 |
| Calcium gluconate (mL) | 10 | 20 | 20 | 20 | 10 | 20 | 30 | 20 | 20 | 20 | 20 | 20 |
| TPN (mL) | 30 | 30 | 30 | 30 | 70 | 30 | 36 | 38 | 30 | 30 | 30 | 30 |
| Albumin 5% (mL) | 250 | 250 | 250 | 250 | 1000 | 700 | 750 | 250 | 250 | 250 | 750 | 500 |

| | | | | | | | | | | | | |
|---|---|---|---|---|---|---|---|---|---|---|---|---|
| **Albumin 25% (mL)** | - | - | - | - | - | - | - | - | 200 | - | 100 | 100 |
| **Creatinine (mg)** | 218 | 218 | 218 | 218 | 218 | 218 | 218 | 218 | 218 | 218 | 218 | 218 |
| **Mannitol (g)** | 5 | 5 | 5 | 2.5 | 4 | 4 | 4 | 2.5 | 2.5 | 2.5 | 0.1 | 25 |
| **Sodium Nitroprusside (mg)** | 5 | 17.5 | 10 | 20 | 22 | - | - | - | - | - | - | - |
| **Velitri (ug)** | 83.3 | - | - | - | - | 164.93 | - | 83.3 | 83.3 | 49.98 | - | 16.66 |
| **8.4% $NaHCO_3$ (mL)** | 30 | 30 | 40 | 60 | 85 | 70 | 30 | - | 20 | 15 | 15 | 35 |
| **Fluid loss replacement (5% albumin) (mL)** | - | 320 | 100 | 700 | - | 450 | - | 250 | 2750 | 1900 | 250 | 2510 |
